# Characterizing Protein Conformational Spaces using Dimensionality Reduction and Algebraic Topology

**DOI:** 10.1101/2021.11.16.468545

**Authors:** Arpita Joshi, Nurit Haspel, Eduardo González

**Author notes:** Arpita Joshi is currently a research fellow at Institute for Systems Biology, Seattle.

## Abstract

Datasets representing the conformational landscapes of protein structures are high dimensional and hence present computational challenges. Efficient and effective dimensionality reduction of these datasets is therefore paramount to our ability to analyze the conformational landscapes of proteins and extract important information regarding protein folding, conformational changes and binding. Representing the structures with fewer attributes that capture the most variance of the data, makes for quicker and precise analysis of these structures. In this work we make use of dimensionality reduction methods for reducing the number of instances and for feature reduction. The reduced dataset that is obtained is then subjected to topological and quantitative analysis. In this step we perform *hierarchical clustering* to obtain different sets of conformation clusters that may correspond to intermediate structures. The structures represented by these conformations are then analyzed by studying their high dimension topological properties to identify truly distinct conformations and holes in the conformational space that may represent high energy barriers. Our results show that the clusters closely follow known experimental results about intermediate structures, as well as binding and folding events.

## 1 Introduction

Characterizing intermediate protein conformations is difficult to do experimentally due to the fleeting nature of these structures. Doing so is necessary for understanding and characterizing protein function and dynamics. This work uses dimensionality reduction and algebraic topology to extract meaningful information inherent in these structures. Protein structure and dynamics are essential to their function. Therefore, understanding the connection between structure, dynamics and function, we can better understand cellular processes involving proteins. The question of how the structure and dynamics of proteins relate to their function has challenged scientists for several decades but still remains open. Conformational search methods aim to characterize the conformational space of proteins in order to find low energy regions corresponding to highly populated or intermediate structures [1], [2], [3]. These intermediate states are transient and therefore hard to detect experimentally. However, they may be essential to understanding dynamic events such as folding, protein-protein interactions and various cellular processes. The potential energy landscape of a protein is often rugged and has a large number of local minima [4]. This makes the conformational exploration especially challenging. The problem becomes even more challenging due to high dimension of the problem. A typical protein can contain several hundreds of amino acids or several thousands of atoms. Therefore, the search space made out of all possible conformations that a protein can assume is large and its enumeration is practically impossible. Existing physics-based computational methods that sample the conformational space of proteins include Molecular Dynamics (MD) [5], Monte Carlo (MC) [6] and their variants, as well as approximate methods based on geometric sampling [2], [7], [8], [9], [10], Elastic Network Modeling [11], [12], [13], normal mode analysis [14], [15], morphing [16], [17] and several other methods.

Even after the conformational space is sampled, it should be filtered and clustered to extract meaningful information. Several clustering methods have been designed for protein conformational space [7], [18], [19]. Most clustering methods for high-dimensional data incorporates metric functions that evaluate the distance between objects in the dataset, or a lower-dimensional representation of these objects, often trying to detect outliers [20].

### 1.1 Problem Statement

The protein folding problem, which aims at predicting the correct protein structure from its sequence, it is an important problem in Biology. The conformational search problem is a related problem that aims at characterizing the conformational space of proteins in order to find minimum energy regions corresponding to highly populated structures and characterize dynamic events such as folding or docking [21], [22]. The potential energy landscape of a protein is immense. A typical protein molecule has hundreds of amino acids and several thousand atoms. Consequently, the number of configurations that a molecule can attain is extremely large and practically impossible to enumerate computationally. This has given rise to a foray of methods that attempt to model the actual pathways that a protein undertakes while transitioning from one conformation to another that ultimately help in elucidating the highly populated conformations generated in the entire process. In this work we present a method to analyze the conformational space of proteins by first reducing the dimensionality of molecular conformations datasets that represent the molecule and then use a number of filtering techniques taken from algebraic topology to identify clusters of intermediate conformations. The molecules used range from small (Oxytocin and Vasopressin) to medium (Cdc42) to large (GroEl). Details about them can be found in the Results section. The main contributions of the paper can be summed up as under:

1. Data representation – Using previous works described in sections 2.2.1 and 2.2.2, we come create a space efficient way to represent molecular data. The datasets in use have just enough instances and attributes that capture the maximum amount of variance.
2. Use versions of hierarchical clustering, depending on the molecule, to sample different conformations of a molecule. Details can be found in section 2.3.1.
3. Use algebraic topology methods to analyze distinctiveness among various clusters of conformations and extract the topological properties of the clusters.

### 1.2 Feature Reduction

It is often helpful to obtain a lower-dimensional representation of the data that preserves as much of the variance in the original data as possible. It is especially useful in protein conformations, since the mutual constraints between atoms in the protein molecule makes the “true” dimensionality of a protein structure much smaller than the number of parameters required to represent a 3 dimensional protein molecule. Existing algorithms for data instance reduction are broadly divided into, incremental, decremental, batch and mixed [23], [24], [25], [26], [27]. The incremental algorithms begin with a null set and data instances are added to it depending on the result of the algorithm. The decremental algorithms, on the contrary, begin with the entire set of instances and depending on the decision offered by the selection algorithms, instances are taken out from the set one had at the beginning. The batch algorithms function in a way that each instance is first analyzed and then a decision is made as to which ones to keep. Mixed algorithms begin with a pre-selected set of instances and the process then continues to figure whether instances should be deleted or added. An evaluation of the age-old techniques of instance reduction is explained in [23].

Linear dimensionality reduction like PCA and its variants may not capture the complex, non-linear nature of protein conformational landscape. Dimensionality reduction techniques are broadly classified based on the solution space they generate, as convex and non-convex [28]. Techniques described in [29] give explicit details of the various well established non-convex methods. These methods are further sub-divided into Full Spectral Techniques, the ones that perform the eigenvector decomposition of a full matrix and Sparse Spectral Techniques, the ones that do the same for a sparse matrix. The latter ones have better time-complexity but these approaches are *local*. They attempt at retaining only the local structure that the sparse portions of the dataset present. On the other hand, Full Spectral Techniques, capture the covariance between all the data instances and form a more thorough representation of the structure as a whole. The Isomap algorithm [30] is a non-linear dimensionality reduction method that falls into the Convex Full Spectral category. It takes as input the distances between points in a high-dimensional observation space, and outputs their coordinates in a low-dimensional embedding that best preserves their intrinsic geodesic distances. The original dimensions of the matrix is *N* × *M*, where N is the number of instances and M is the size of each instance. The output matrix is of size *N* ×*m*, where *m* << *M*. In a previous work [2] we have shown that for protein datasets, Isomap produces much better results, due to protein conformational changes being non-linear and complex. Despite its advantage in efficient representation of molecular data [31], [32], Isomap is computationally expensive, especially with very large, multi-dimensional datasets. To overcome this, we have implemented our version of the Mode-III of Isomap. Improvements over Isomap are presented in [33], [34]. A similar approach is adopted for this work but in a way that is more suited for protein data.

### 1.3 Algebraic Topology

The conformational space of proteins is highly complex and high dimensional. Obtaining a full, analytical description of it is essentially computationally impossible. Different methods for characterizing the conformational landscape try to obtain a coarser, more approximate description of the landscape while still capturing its global, essential characteristics while preserving important features. Below we give a brief survey of the tools used in this paper.

#### 1.3.1 Persistent Homology

Persistent homology is an algebraic topological tool for computing features of a space at (essentially) different spatial resolutions [19]. To find the persistent homology of a space *X*, presented as a data set, we first assign a *simplicial complex*. Moreover, using a distance function on the underlying space, we can build a *filtration* of the simplicial complex, that is, a nested sequence of complexes. We follow the notations in [35].

##### 1.3.1.1 Simplicial Complexes

A simplicial complex *K* is given by the following datum:

- A set Ko of vertices or 0-simplices. We will also use the notation *K*_0_ =*Z* for consistency with the literature.
- For each *i* ≥ 1, a set *K_i_* of *i*-simplices *σ* = [*z*_0_*z*_1_…*z_i_*], where *Z_j_* ∈ *Z*.
- Each *i*-simplex has *i* + 1 *faces σ_j_*, *j* = 0,…, *i* obtained by deleting the *j*-th element in the list. We require that the faces *σ_j_* are in *K*_*i*–1_.

We think of 0-simplices as vertices, 1-simplices as edges, 2-simplices as triangular faces, and 3-simplices as tetrahedra, etc.

##### 1.3.1.2 Homology and Betti numbers

For a simplicial complex *K* as above, we associate the group of *i*-chains *C_i_* with coefficients on a ring *R* (we will take *R* =**Z**_2_ to avoid issues with orientations) as the group generated by the elements of *K_i_*, that is the set of formal sums Σ*r_s_σ_s_*, *σ_s_* ∈ *K_i_*, *r_s_* ∈ *R*. The boundary map *∂_i_*: *C_i_* → *C_i–1_* takes an *i*-simplex *σ* = [*z*_0_,…, *z_i_*] ∈ *K_i_* to the formal alternating sum Σ(—1)^*j*^ [*z*_0_,…, *ẑ_j_*,…, *z_i_*], and it is extended by linearity over *C_i_*. This map satisfies *∂_i_*○ *∂*_*i*+1_ = 0. That is, the *boundaries* given by the image *B_i_* = im(*∂*_*i*+1_) ⊂ *C_i_* lies in the subgroup of *cycles*, given by the kernel *Z_i_* = ker(*∂_i_*) ⊂ *C_i_*. This makes C_●_ a chain complex. The *i*-th homology *H_i_*(*K*) is defined as the quotient *Z_i_*/*B_i_*, this is a module over R.

The *k*-*th Betti number Betti_k_*(*K*) of the complex is the rank of the *k*-th homology of the complex *H_k_*(*K*) as an *R*-module. Roughly speaking, this gives a count of the number of *k*-dimensional holes in the complex. In particular, *Betti_0_*(*K*) is the number of connected components of *K*. For instance, a *k*-dimensional sphere, has all Betti numbers equal to zero except for *Betti*_0_ = *Betti_k_* = 1. For a topological space *X*, approximated by a complex K_X_, its homology *H_i_*(*X*) = *H_i_*(*K_X_*) and associated Betti numbers are classical invariants. Figure-1 has some examples, for instance a torus has one connected component so *Betti_i_* =1, two non-homologous one-dimensional cycles - one equatorial and one meridional, so *Betti*_1_=2 and a single 2-dimensional hole enclosed within the surface, *Betti*_2_=1. Similarly, for a circle, *Betti*_0_= *Betti*_1_= 1, and for a point *Betti*_0_= 1.

**Fig. 1.**
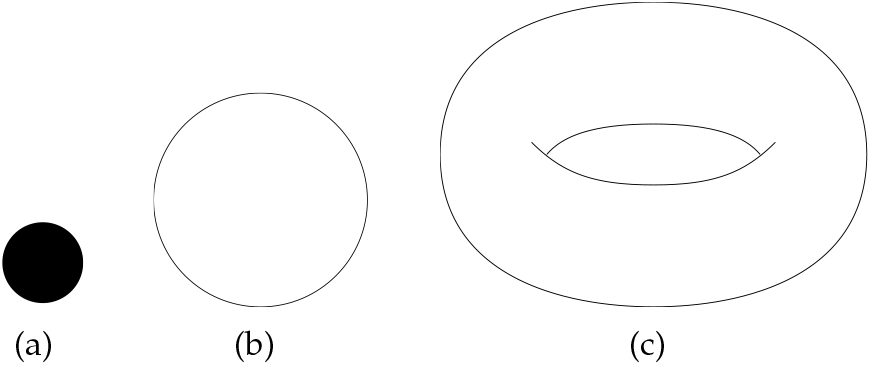
Representation of Betti Numbers as holes: (a)A dot – 1 connected component, (b) A circle – 1 connected component and 1 loop, (c)A torus – 1 connected component, 2 loops and a 2-dimensional face.

##### 1.3.1.3 Data Sets and Persistent Homology

An important example for this paper consists of the simplicial complex associated to a data set or point cloud. Let *Z* be a set of points in an Euclidean space 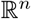. To this data, and to any radius *ϵ* > 0 we can associate a simplicial complex *K_ϵ_* as follows. The 0-simplices consist of the points {*z*} in the data set themselves. A set of two points {*z*_0_, *z*_1_} is declared to be a 1-simplex if the *ϵ*-balls centred at *z_i_* intersect, that is if *B_ϵ_*(*z*_1_)⋂*B_ϵ_*(*z*_1_) = Ø. Analogously the *k*-simplices will be defined by those sets of *k* + 1 points {*z*_0_,…, *z_k_*} for which the intersection *B_ϵ_*(*z*_1_) ⋂…⋂*B_ϵ_*(*z_k_*) = Ø is not empty. Certainly for small *ϵ* the complex only contains 0-complexes and for large *ϵ* the complex yields the *degenerate* complex for which all sets of *k* + 1 points define a *k*-simplex since all the points will be inside a single ball. Varying the ϵ yields a filtration of this complex. By a filtration of a simplicial complex *K* we mean a collection of sub-complexes 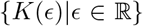 of *K* such that *K*(*t*) ⊂ *K*(*s*) whenever *t* ≤ *s*. A filtration yields *Persistent Homology*, an *ϵ* dependent homology theory H_ϵ_ and hence a family of Betti numbers which depend on *ϵ*. This is what its called *persistent bar codes*. For all purposes, we will interpret the radius *ϵ* as a time parameter and we will sometimes denote it as *t*. We are interested in the study of how the topological invariants of a dataset varies with *t* and which Betti numbers and generators of homology persist with this variation. Several computational tools to compute Persistent Homology and associated bar codes, such as JavaPlex [36] and the Topological Data Analysis TDAToolBox [37] for R and [38] for Python were used. In general one has to apply dimensionality reduction methods to effectively compute any topological analysis.

In this work, on the embeddings produced by dimensionality reduction, we perform both hierarchical clustering and compute its persistent homology. The Betti_0_ of the embeddings produces the same number as the number of relevant clusters in hierarchical clustering. The results are presented in the next section. Furthermore, these clusters are then subjected to further topological analysis. Each of these clusters (of a molecule) is a connected component of the embedding. Consequently, the Betti_0_ of each of these is 1. We observe *Betti_1_* and *Betti*_2_ (loops and voids respectively) of each of these clusters to find out about the topological properties of the distinct clusters.

Algebraic topology in Bioinformatics: Algebraic topology approaches have recently been explored in the context of sampling biological data. The work cited in [39] describes the use of topological signatures, which the authors call evolutionary homology (EH) barcodes, reveal the topology-function relationship of the network and thus give rise to the quantitative analysis of nodal properties. The proposed EH is applied to protein residue networks for protein thermal fluctuation analysis, rendering accurate B-factor prediction for a number of proteins. A thorough review of the emerging applications of topological methods to genomics is presented in [40]. An insightful piece presenting concisely the ramifications of these approaches are elucidated in [41]. Algebraic topology and persistent homology were also used for protein-protein interaction [42] and protein conformational dynamics [19]. Previously, we were able to detect intermediate structures in protein conformational samples using algebraic topology and hierarchical clustering [3], but we did not explore higher order betti numbers that may give us more information about the conformational space. In this contribution we fill the gap by analyzing the conformational spaces of multiple proteins of different sizes using persistent homology, clustering and dimensionality reduction techniques. Our results match experimental data when it is available.

## 2 Methods

### 2.1 Generation of Data

The molecules chosen with in this work are diverse and very different in their function and dynamics. The conformational spaces for oxytocin, vasopressin, hGalanin, pGalanin and CDC-42 were sampled using Molecular Dynamics(MD) simulations. All systems were represented at full atomic resolution. All the above proteins except CDC-42 are shorter peptides with no crystal structures. The initial linear peptide was modeled using the Chimera software [43] as an idealized helix. Hydrogens and solvent were added using the AMBER software package [5]. Simulations were performed in constant volume (NVT) in an orthorhombic box using the TIP3 solvent model [44] to simulate infinite dilution.

Periodic boundary conditions were applied using the nearest image convention. The overall charge of the system was kept neutral for the use of particle mesh Ewald summation to calculate electrostatic charges [45]. The simulations were carried out with the NAMD package [46] using the AMBER ff03 force field [47]. To sample the conformational spaces of the peptides we used a simulated annealing based search [6]. For CDC-42, we used long MD simulations as described in [48]. The GroEL molecule is larger and cannot be easily sampled using MD simulations. The conformations were sampled using a Monte Carlo (MC) [6] based conformational pathway search. The protocol is detailed in [3]. The search begins with the PDB (Protein Data Bank) format of one conformational extreme and expands following a biased Rapidly-expanding Random Tree (RRT) algorithm to simulate the pathway that can be undertaken to reach the goal conformation [49]. At every iteration a parent protein conformation is chosen from pool of new conformations, the one selected is the one which has its energy below a threshold ^1^. The new conformation is added to the pool^2^ if its RMSD (Root Mean Square Deviation) is closer to the goal.

### 2.2 Dimensionality Reduction

#### 2.2.1 Data Instance Reduction

The first step is to perform dimensionality reduction on the data, to obtain a reduced representation. We use a data instance reduction method to decide the information content offered by each data instance. We first apply spherical PCA as described in [50], [51], [52]. The input is a data matrix and the number of dimensions (principal components) desired of all the data instances. The output is a lower dimensional projection of the data. We used two or three dimension projections, in order to better visualize structurally significant data. It is observed that over eighty percent of the variance is explained by the first three principal components in all the types of data we used, although it is not always the case in other domains. Our algorithm removes data instances that are deemed not to contribute to the variance in the data. More details about the algorithm itself and the results it produced can be found in [53], [54]. The result of this step produces a non-redundant representation of the dataset which now has fewer instances but its important properties that account for maximum variance present in it are preserved.

#### 2.2.2 Low-Dimensional Embedding

The next step is to obtain an embedding of the conformations in the reduced dataset in a lower dimension. We used a parallelized version of the Isomap algorithm that produces the same results and works much faster [55]. The algorithm can be used to produce an embedding in as many dimensions as the attributes of the dataset, we pick the first three dimensions to work with because they capture over 80% of the variance inherent in the data.

### 2.3 Topological Analysis

After obtaining a reduced embedding that represents the dataset, the next step is to slot the data to sample different conformations inherent in these embeddings. To ascertain the number of conformations that can be extracted, we use two methods, namely, hierarchical clustering and persistent homology. Persistent Homology is the next step in the process but here it was used as a proof of validation, as both, hierarchical clustering and persistent homology revealed the same structural complexities in each dataset. We used the R package TDA’s (Topological Data Analysis) function *calculate*_*homology* [56] and the TDAToolBox [37] to generate the topological analysis.

#### 2.3.1 Hierarchical Clustering

Hierarchical Clustering is a known method for identifying similar groups in a dataset. Built-in function in R, *hclust* is used for the purpose. We chose hierarchical clustering over k-means clustering to prevent having to pre-declare the number of clusters sought, and present the results coherently in the form of a dendrogram. Depending on how similar or dissimilar the data instances are, hierarchical clustering can be divided into two categories:

- Agglomerative Hierarchical Clustering: It works in a bottom-up fashion. Each data instance is considered a single element cluster to begin with. At each step of the algorithm, two most similar clusters are combined into one. The process continues until all instances have been combined into one big cluster containing all the data instances.
- Divisive Hierarchical Clustering: This method works complementary to the previous one. Here, all the data instances are considered a point of one big cluster at the beginning. At each iteration of the process, the most heterogeneous cluster is divided into two. The process continues until all the data instances are a single element cluster.

Agglomerative form of hierarchical clustering is more suited for more heterogeneous data that has a large number of small clusters to identify. On the other hand, divisive clustering as expected, is better at isolating big clusters. We use both these approaches depending on which molecule we are analyzing. The molecular datasets that are known to undergo large scale conformational changes like Calmodulin, Adenylate kinase and GroEl were subjected to agglomerative hierarchical clustering and the others to the divisive one. To isolate the clusters, we plot the results and the dendrogram tree suggests where it can be cut for major clusters. The function, *cutree* serves the purpose and the function *table* computes the size of each of these clusters. Then the clusters with at least 100 instances are picked. For example, in Figure-2 the width of the clusters is suggestive of their sizes, if the cutree function is called with an argument grater than 2, one of the two clusters splits in a way such that one child cluster has fewer than 100 points.

**Fig. 2.**
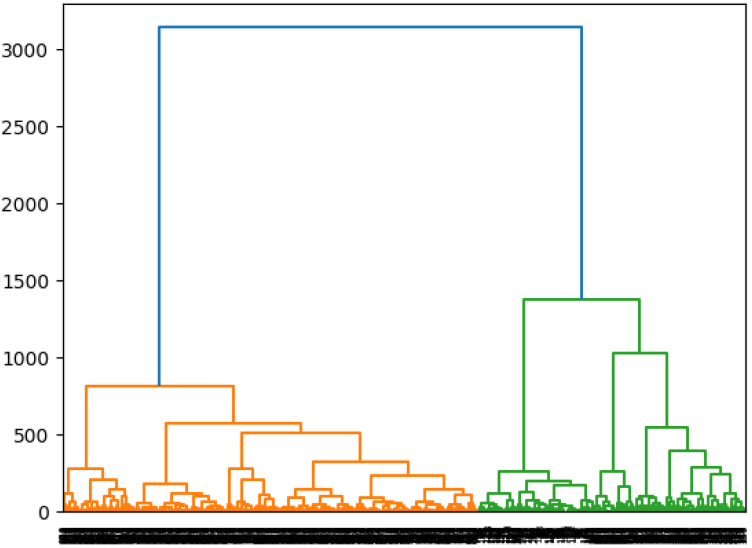
Clustering hierarchy for the inactive GDP-bound conformation of CDC-42. The conformational space breaks into two clearly defined clusters.

The entire workflow can be summed up as under:

1. Obtain a subset of the dataset generated by MD simulations using the PCA based data instance reduction.
2. Obtain the low dimensional embedding by performing Isomap on this subset.
3. Perform hierarchical clustering on this embedding and compare to the *Betti_0_* value for estimating the number of clusters.
4. Compute *Betti*_1_ and *Betti*_2_ of the clusters to establish the different topological properties of distinct clusters.

## 3 Results and Discussion

In this work we chose a number of medium to large proteins with different conformational dynamics. As mentioned earlier, to generate clusters from the reduced dimension embedding various methods were used depending on the molecular dataset. A thorough analysis for the various molecules is as under.

### 3.1 CDC-42

Cell Division Control-42 is a GTP (Guanosine Triphosphate) binding protein [57]. Human CDC-42 is a small molecule with 191 amino acids. It belongs to the *Rho* protein family, which regulates signaling pathways that control a wide range of cellular functions, for example, cell migration and cell cycle progression. It switches between cycles an active GTP-bound state and an inactive GDP (Guanosine Diphosphate)-bound state. Recently, CDC-42 has been shown to actively assist in cancer progression and metastasis. Several studies have established the basis for this and hypothesized about the underlying mechanisms [58], [59]. The data for all of the molecules was procured using Molecular Dynamics trajectories from a recent work by Haspel et al. [48].

#### Inactive form (GDP bound)

Both forms of hierarchical clustering produced the same number of clusters here, shown in Figure-2. There were two significant clusters. We call a cluster significant if it has at least 100 data instances. Figure-3 shows the projection obtained by the feature reduction algorithm, where each cluster is shown in different color. To visualize the topology of the conformational landscape we used the kernel density estimation method and persistence diagram as described in the TDAToolBox package [60], Figure-4(a) shows the density estimates over the embedding, highlighting two peaks, Figure-4(b) shows the persistence diagram for the conformational space of this molecule. At the given *ϵ* as defined and elaborated in section 1.3), here *Betti*_0_ = 2, tantamount to two relevant clusters of this molecule. One conformation representative of each of the significant clusters is shown in Figure-5 (a) and (b). Both these conformations are selected as the data point closest to the centroid of the two clusters. The PDB (Protein Data Bank) representations shown in Figure-3 were obtained using VMD (Visual Molecular Dynamics) [61]. Figure 5(c) shows a superimposition of the two clusters with the regions associated with binding and activation highlighted in color: The Switch-I region (residues 25-37) in blue, the Switch-II region (residues 57-75) in orange and the insert region (residues 122-135) in red. It can be shown that these three regions represent the highest variability, consistent with the fact that they are involved with binding and activation [62], [63]. These two clusters are then compared to each other to deduce conformational information.

**Fig. 3.**
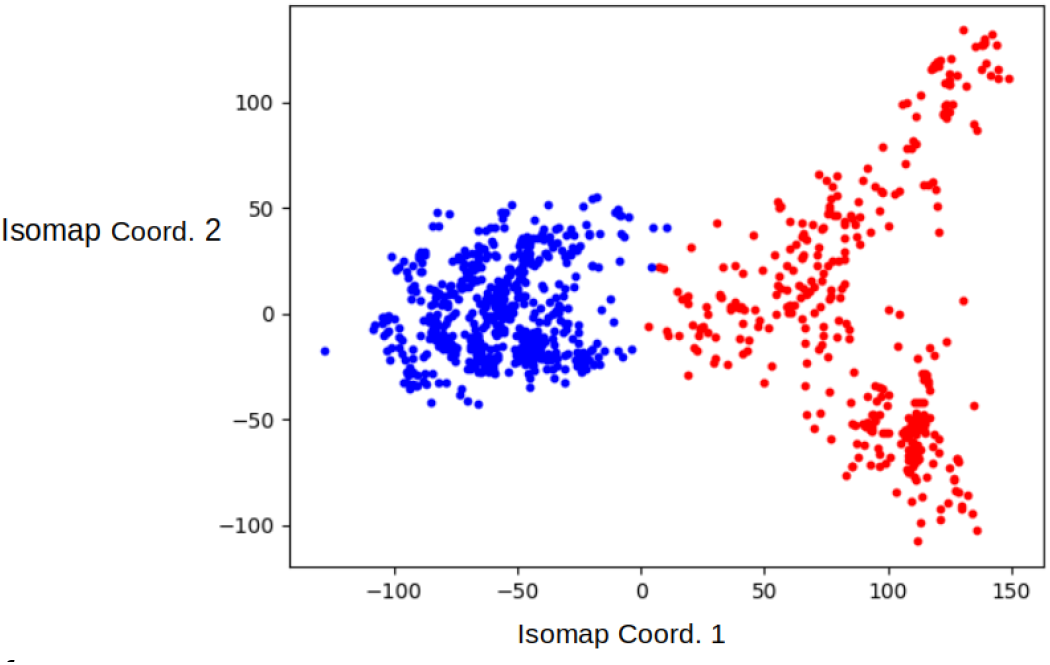
Embedding of the two clusters of the inactive Cdc42 (PDB:4did). Each cluster is highlighted in a different color.

**Fig. 4.**
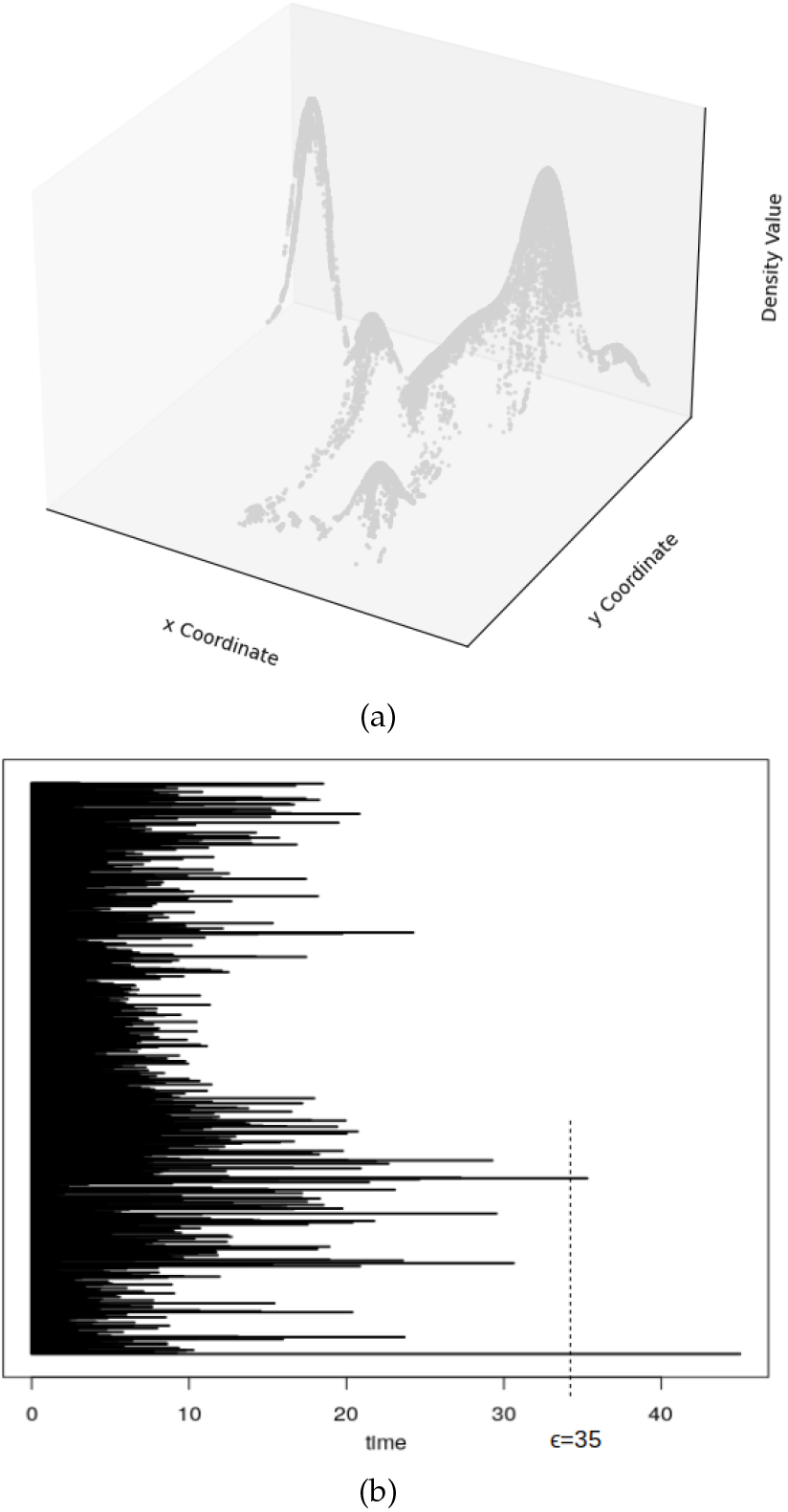
Density Measurements for Inactive CDC42 (4did): (a) Density Estimate over 3-D space, (b) Persistence barcodes for two clusters persist at the given level of filtration for conformational space.

**Fig. 5.**
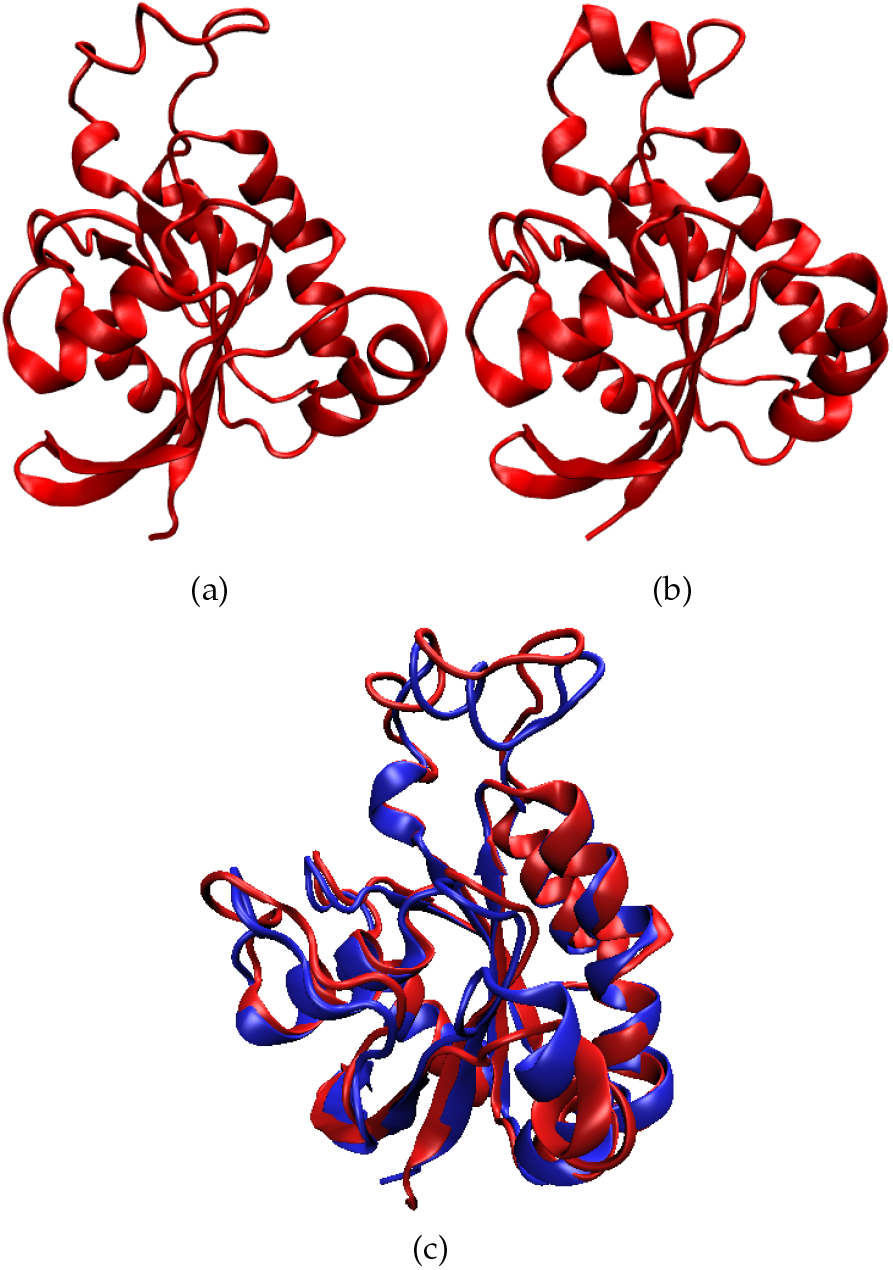
(a), (b) Two conformations of the inactive form of CDC-42 (PDB 4did)(c) A superimposition of the two clusters, the regions with maximum difference stand out at the top of the figure.

Next, we examined the topological features of each cluster separately. As mentioned earlier, all of the clusters (in all the molecules) are one connected component of a larger structure, so *Betti*_0_ of all of them is 1. A quantitative representation of the persistence barcodes for the two clusters is shown in Figure-6. It is another way of visualizing persistent homology. Each feature is depicted by a single point with the horizontal axis representing feature birth and the vertical axis representing feature death. The line *y* = *x* is included as a reference. Since feature birth always precedes feature death (*x* < *y*), all points in a persistence diagram lie above the reference line. Just like topological barcodes, feature dimension is coded as the point’s color, the red dots represent dimension 0 (*Betti*_0_), the blue dots, dimension 1 (*Betti*_1_) and the green dots represent dimension 2 (*Betti*_2_). In Figure-6, we see a number of 0-cycles (red dots); it is difficult to differentiate between most individual 0-cycles due to all of them being a part of the same connected component. However, there is a number of loops (blue dots); in Figure-6 (b) three of which are significant and persist in the entirety of the conformational space, there is also a green dot (a void) that persists. This represents the bigger, more branched and perforated cluster of Figure-3, the other cluster of this figure is smaller and structurally not as complex, as is also evident from its *Betti*_1_ distribution in Figure-6(a), none of the loops in this conformation are significant enough to span the entire conformational space. The significant thing to note here is that both of these conformations have different distributions of the higher Betti number at the same level of filtration (*ϵ*) which means that conformations represented by these clusters are distinctively different.

**Fig. 6.**
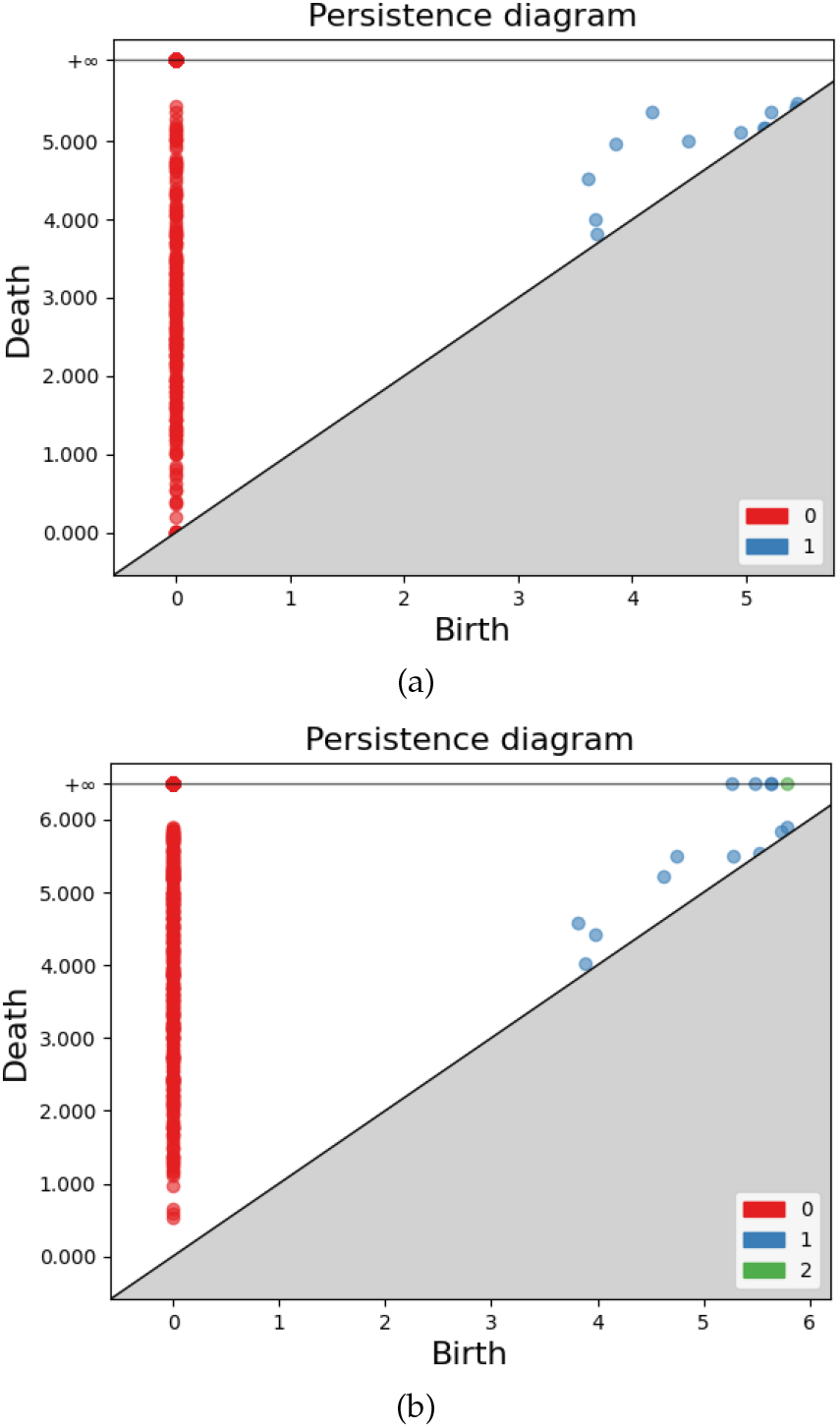
Quantitative representation of the persistence barcodes for the two clusters of the inactive (GDP bound) state of CDC-42: Red dots are Betti_0_ and the blue dots represent Betti_1_.

#### Active form

This active (GTP bound) form of CDC-42 was subjected to divisive form of hierarchical clustering that that indicated that the molecule has only one significant cluster. The clustering hierarchy, isomap embedding, persistence diagram and the density estimates for this form of CDC-42 can be found respectively in Figure-7, all of which corroborate the existence of only one conformation. Figure-7 (a) clearly shows that there is only one significant cluster, the right part of the dashed line is a number of data points much less than 100 and hence is not significant.

**Fig. 7.**
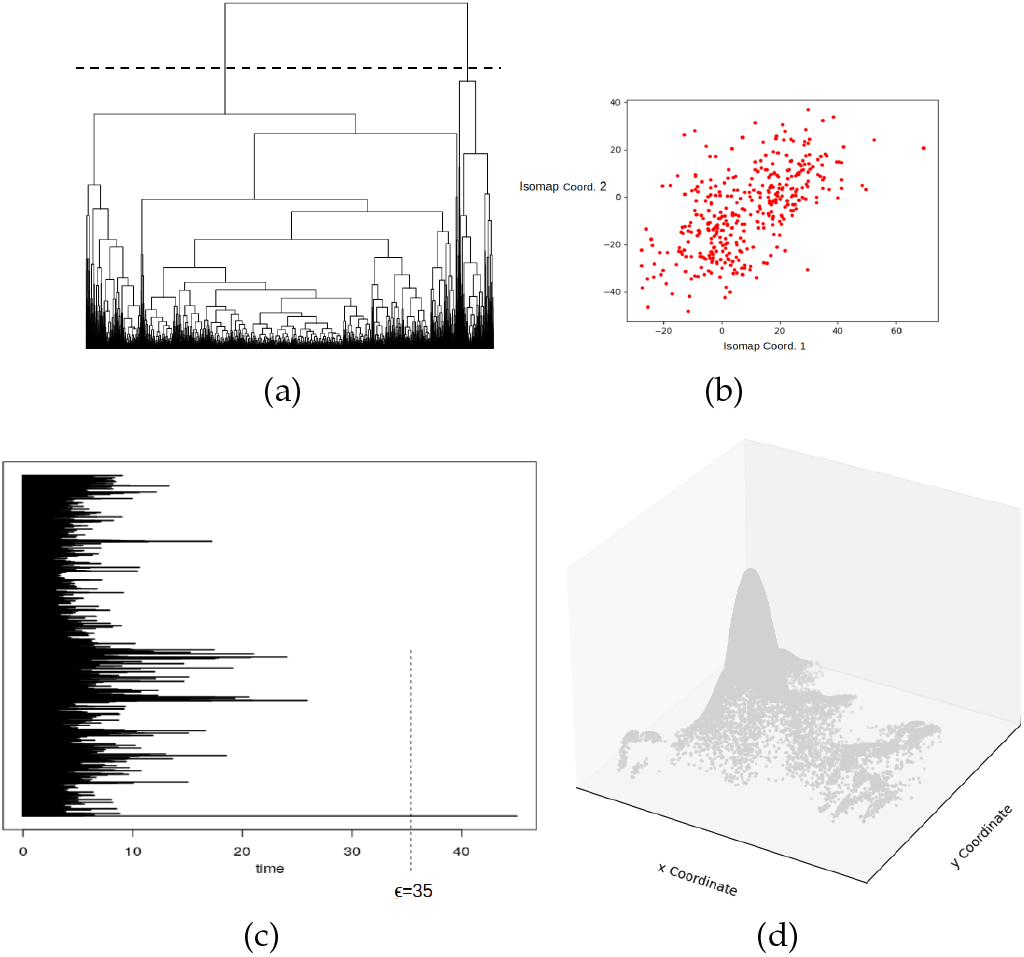
Active form of CDC-42-(PDB 4js0): (a)- Divisive Clustering hierarchy, (b)- Isomap Embedding, (c)- Persistence diagram with *ϵ* = 35, (d)- Density estimates

### 3.2 oxytocin and Vasopressin

Oxytocin and Vasopressin are both small peptide hormones, both with 9 amino acids [64]. They differ only in two positions in their amino acid sequence, as shown under:

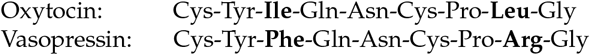

These are smaller peptides that do not have one stable structure. So, intuitively, divisive hierarchical clustering works better to sample conformations here. Nonetheless, for oxytocin we performed agglomerative clustering and obtained a number of clusters but they were very small and exhibited the same higher Betti numbersThe first three Betti numbers of these five clusters are in Table-1. These numbers indicate that there really are just two different clusters here: a result that was obtained when divisive form of hierarchical clustering was performed which resulted in just two significant clusters. This shows that the choice of the type of hierarchical clustering is critical to the study of conformational landscape. The hierarchy produced by divisive clustering is shown in Figure-8(a). As seen in Figure-8 (b), the kernel density estimations for Oxytocin also resulted in two clusters. Figure-8 (c) the persistence diagram for the entire conformational space shows three red dots that ultimately represent two clusters - one big and one small, smaller of which is made of two even smaller clusters that merged during the process. The persistence diagrams for the two clusters of Oxytocin and the one in Vasopressin are shown in Figure-9. The first cluster had two loops and no voids. The second cluster on the other hand had three loops and a void. For Vasopressin, there was only one significant cluster produced by divisive hierarchical clustering. The only cluster in Vasopressin is close in its topological structure with the known Oxytocin conformation which was expected given the similarity in the sequences of the amino acids of the two molecules. The conformations representative of the two clusters of Oxytocin and the one of Vasopressin are shown in Figure-10.

**Fig. 8.**
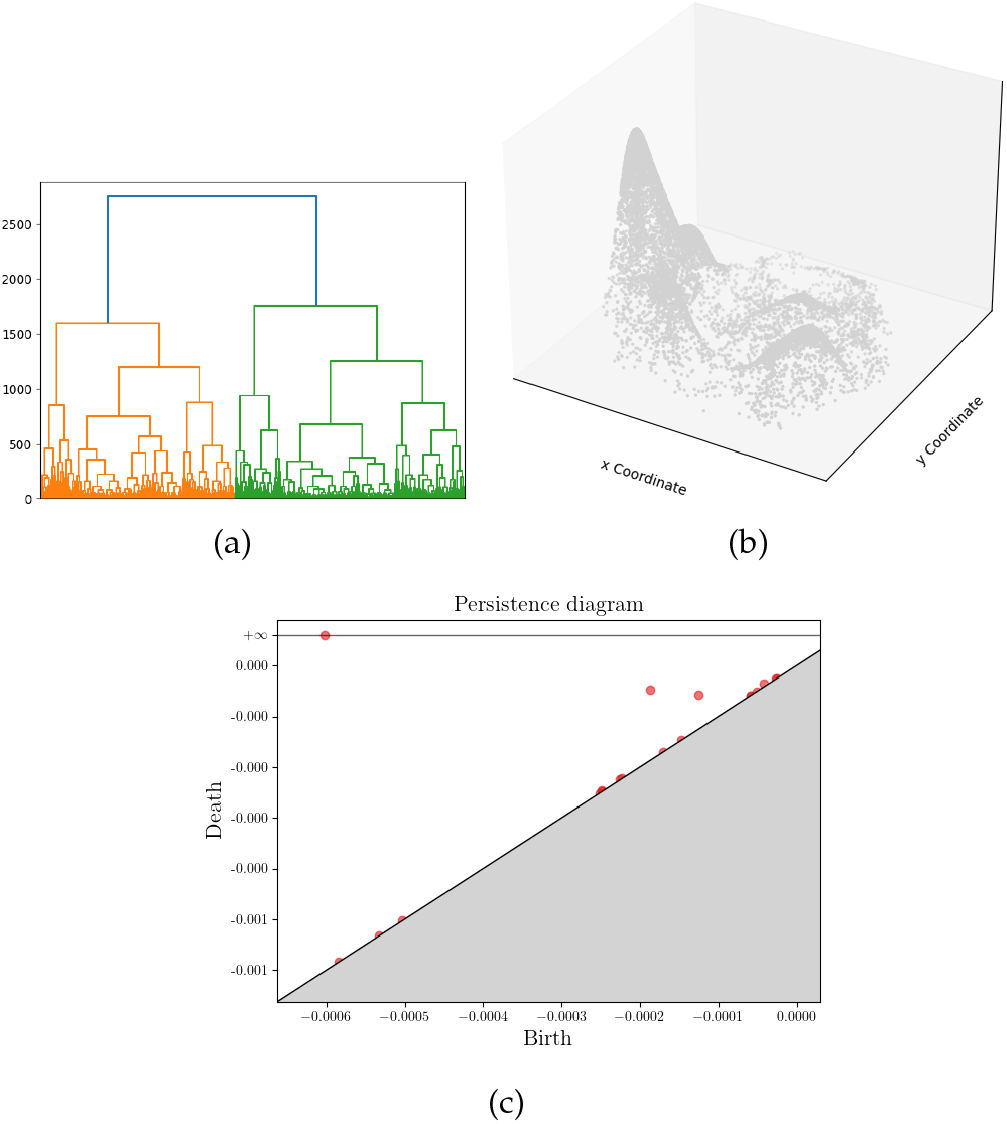
Oxytocin: (a)- Divisive Clustering hierarchy, (b)- Density Estimate, (c)- the persistence diagram, showing one persistent cluster and two others (on the top right) that soon merge into the main one.

**Fig. 9.**
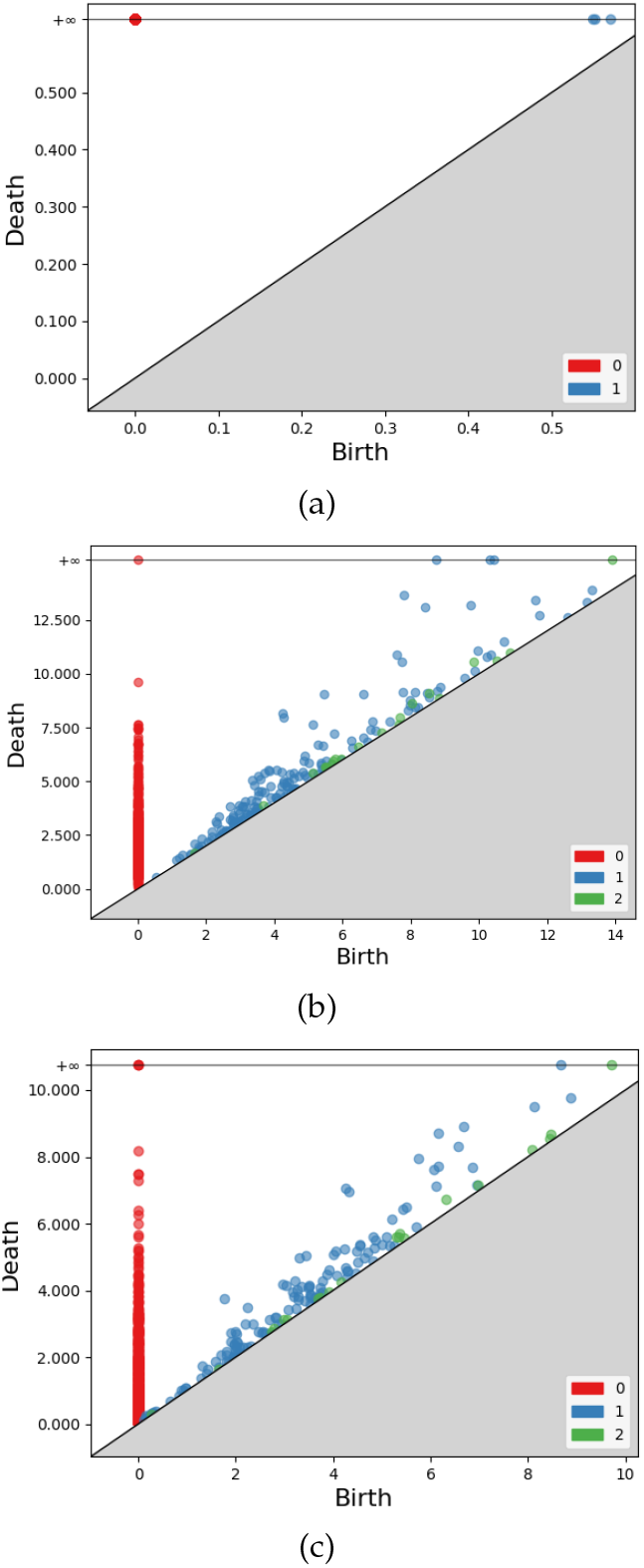
Persistence Diagrams of: (a)- Oxytocin, first cluster, (b)- Oxytocin, second cluster, (c) Vasopressin.

**Fig. 10.**
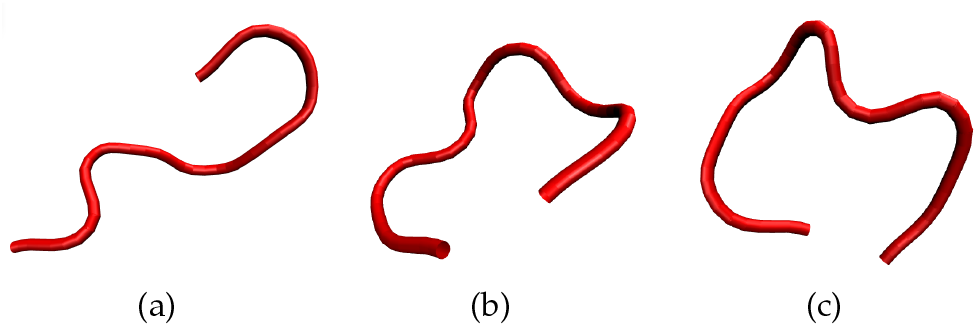
PDB Conformations of: (a)- Oxytocin, first cluster, (b)- Oxytocin, second cluster, (c) Vasopressin.

**TABLE 1.** Betti Numbers for clusters of Oxytocin produced by Agglomerative Hierarchical Clustering.

| Cluster No. | $Betti_0$ | $Betti_1$ | $Betti_2$ |
| --- | --- | --- | --- |
| 1 | 1 | 1 | 0 |
| 2 | 1 | 1 | 0 |
| 3 | 1 | 1 | 0 |
| 4 | 1 | 4 | 0 |
| 5 | 1 | 0 | 0 |

### 3.3 Human and Porcine Galanin

Galanin is a neuropeptide which is encoded by the GAL gene. The gene is expressed in the parts of central nervous system, and gut of humans and some other mammals [65]. Porcine Galanin (pGalanin) and Human Galanin (hGalanin) are two variants of Galanin that differ by 5 out of their amino acids. pGalanin contains 29 amino acids and hGalanin has 30. Both of these molecules also do not have one stable 3D structure due to their small size. Divisive hierarchical clustering on hGalanin resulted in two major clusters (shown in Figure-11) and that of pGalanin produced only one cluster.

**Fig. 11.**
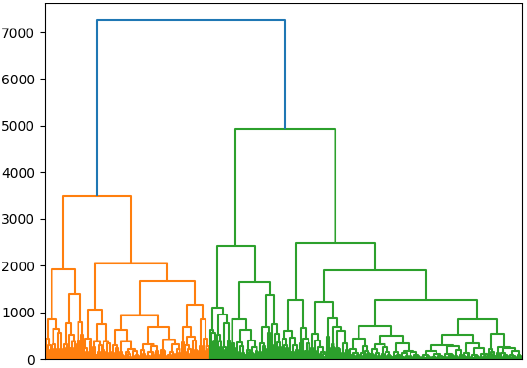
Divisive clustering hierarchy of Human Galanin.

Topologically, the two clusters of hGalanin are quite different from each other, shown in Figures-12(a) and (b), which is in agreement with experimental results produced by Holst et al [66]. The persistence diagram for pGalanin is shown in Figure-12(c). As is evident from the persistence diagrams, the second cluster of hGalanin and the pGalanin cluster are very similar. The study of hGalanin in [66] also shows that there are two molecular forms of hGalanin, one of 30 and another of 19 amino acids. The larger of the two peptides has a sequence which is identical to that of pGalanin except for the following substitutions: Val16, Asn17, Asn26, Thr29 and Ser30. The PDB structures of these clusters is shown in Figure-13, further corroborating that there are two conformations for hGalanin and one of them is structurally similar to that of pGalanin.

**Fig. 12.**
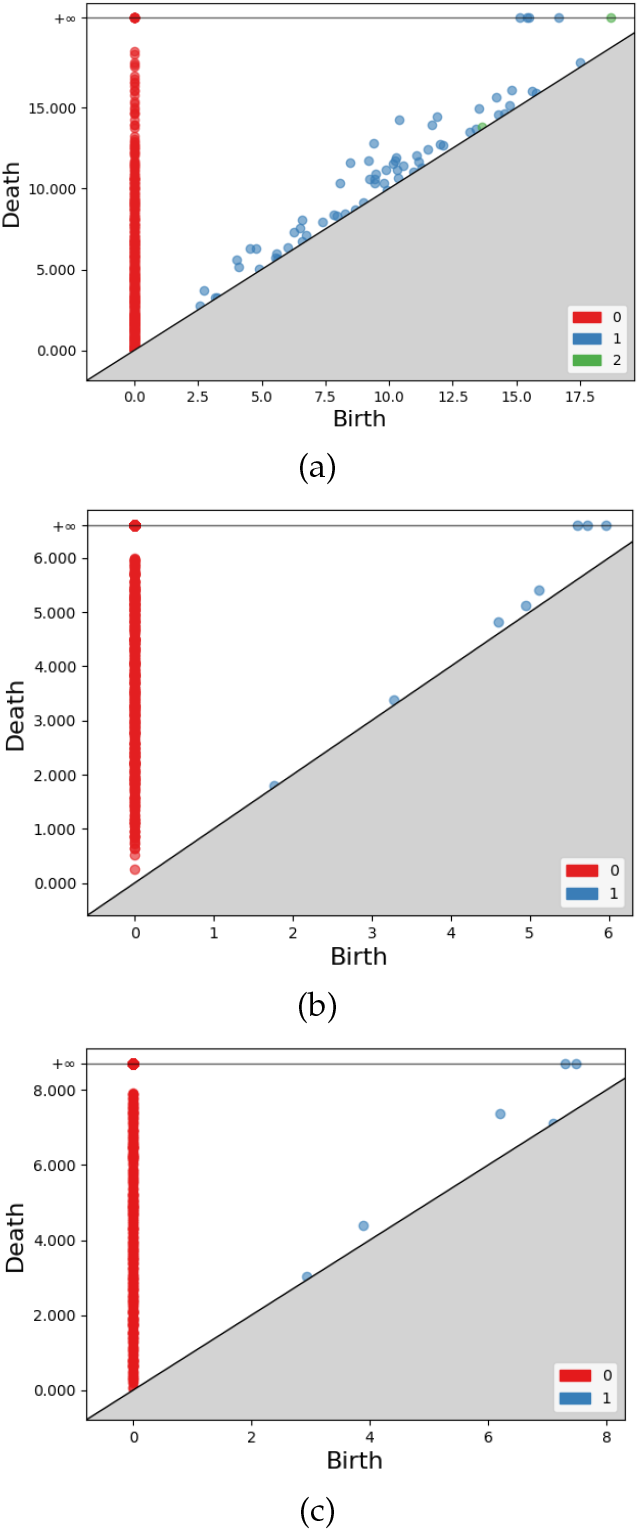
Persistence diagrams for the (a)- first cluster of hGalanin, (b)- second cluster of hGalanin, (c)- pGalanin cluster.

**Fig. 13.**
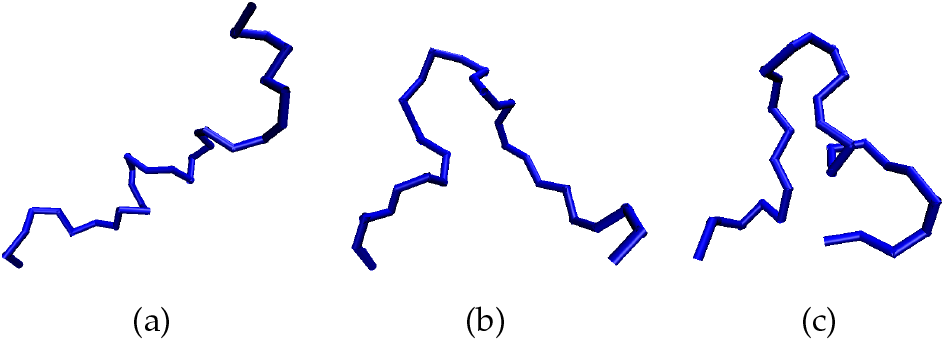
Conformations of Galanins: (a-b)- Conformations representative of the two clusters of hGalanin, (c) Conformation representative of the single pGalanin cluster

### 3.4 GroEL

GroEl is a chaperonin protein which is needed by many other proteins for their proper folding. Structurally, GroEL is a dual-ringed tetradecamer, with both the *cis* and *trans* rings consisting of seven subunits each. The conformational changes that occur within the central cavity of GroEL cause for the inside of GroEL to become hydrophilic, rather than hydrophobic, and is likely what facilitates protein folding. GroEL requires the co-chaperonin protein complex, GroES. Binding of substrate protein, in addition to binding of ATP, induces an extensive conformational change that allows association of the binary complex with GroES. It is the heaviest molecule in the study with 524 amino acids. It is known to undergo large scale conformational changes and hence agglomerative form of hierarchical clustering was used here. It helped in identifying six different clusters (as is corroborated by a previous work [3]), isomap embedding and clustering hierarchy of which is shown in Figure-14 (a) and (b) respectively. This is the most diverse molecule in the study and has a complex more perforated conformational space, and hence the isomap embedding is shown in three dimensions to highlight the different clusters. The conformations representative of each of these clusters are shown in Figure-15. Looking at the clusters, one can see the opening and closing along the hinge. The persistence diagrams for the six individual clusters can be found in Figure-16.

**Fig. 14.**
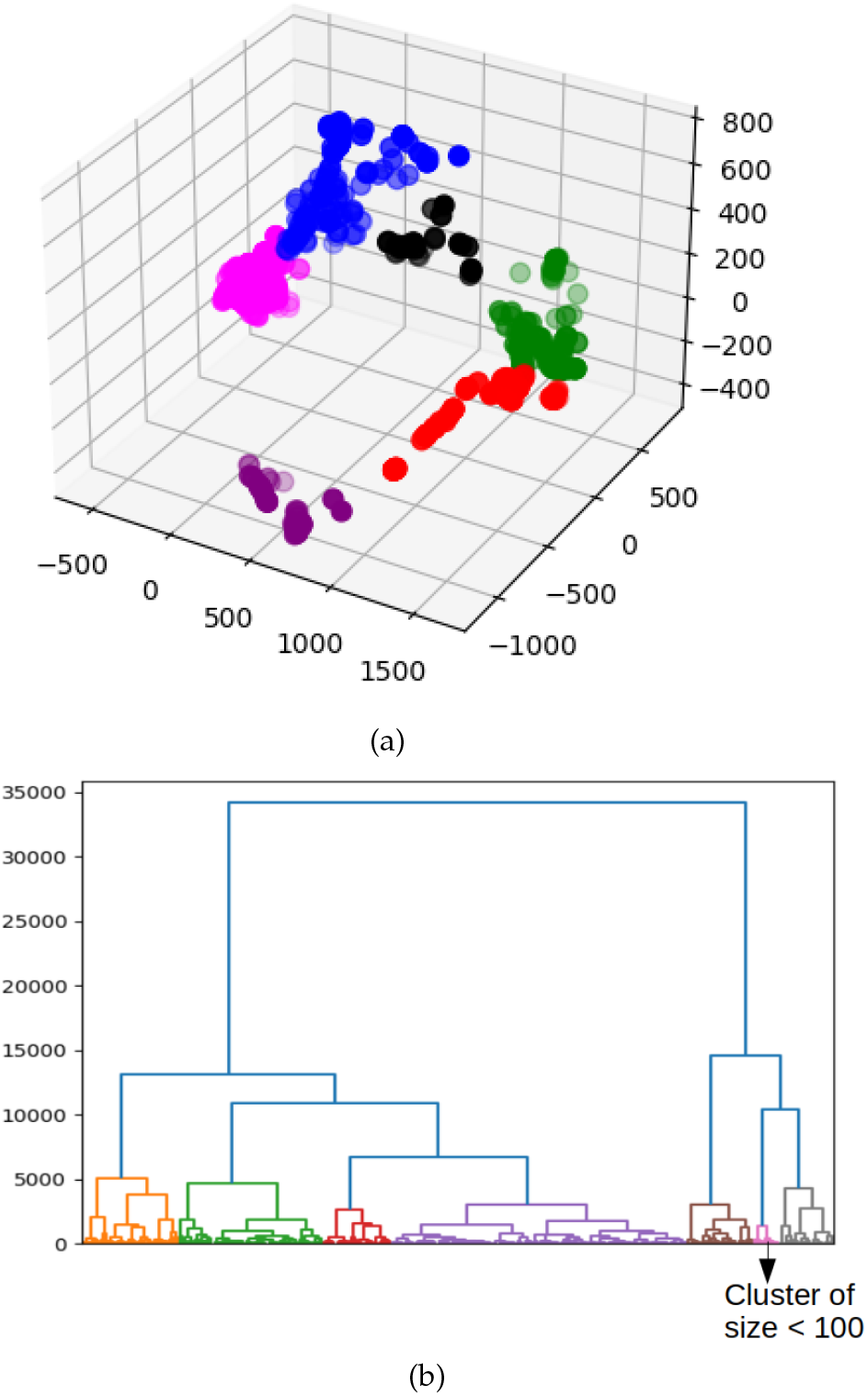
(a)- Embedding of the six clusters of GroEL. Each cluster is highlighted in a different color. X, Y, Z axes are the respective Isomap coordinates, (b)- Agglomerative clustering hierarchy, the insignificant cluster was taken off the analysis.

**Fig. 15.**
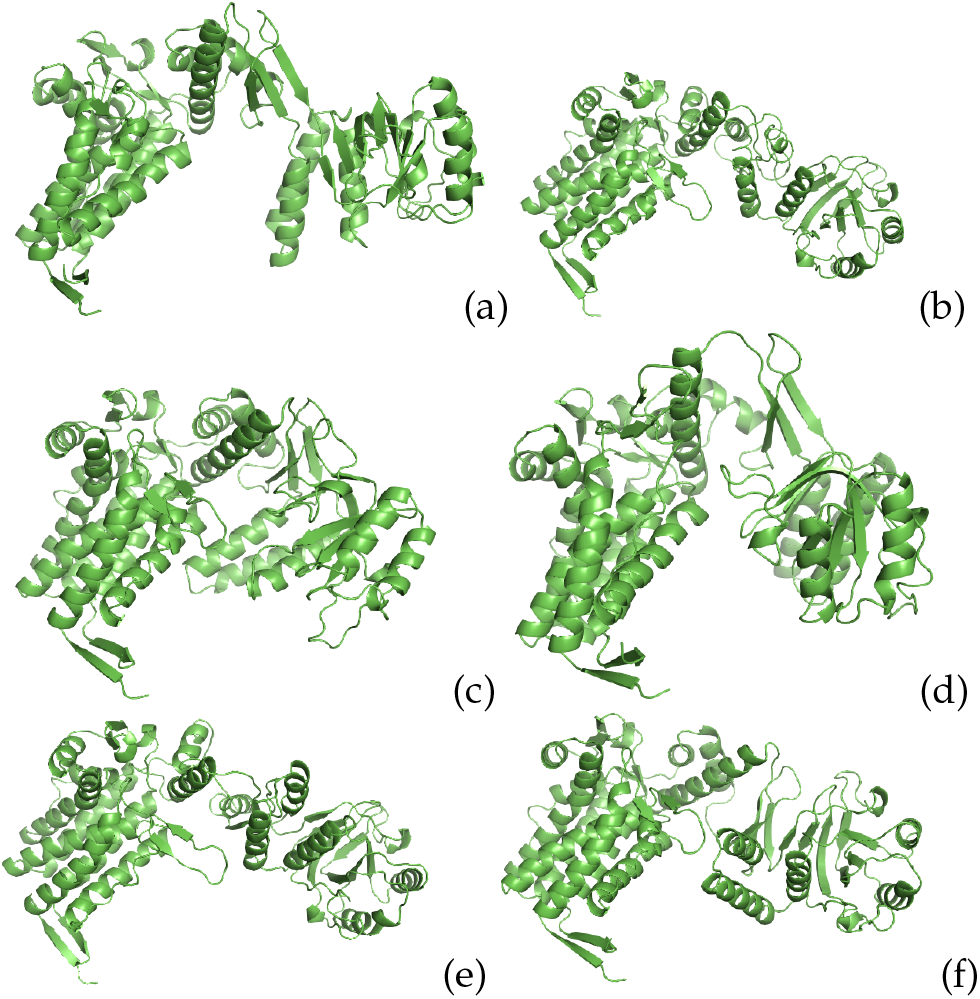
Representatives of the six clusters obtained for the GroEL monomer.

**Fig. 16.**
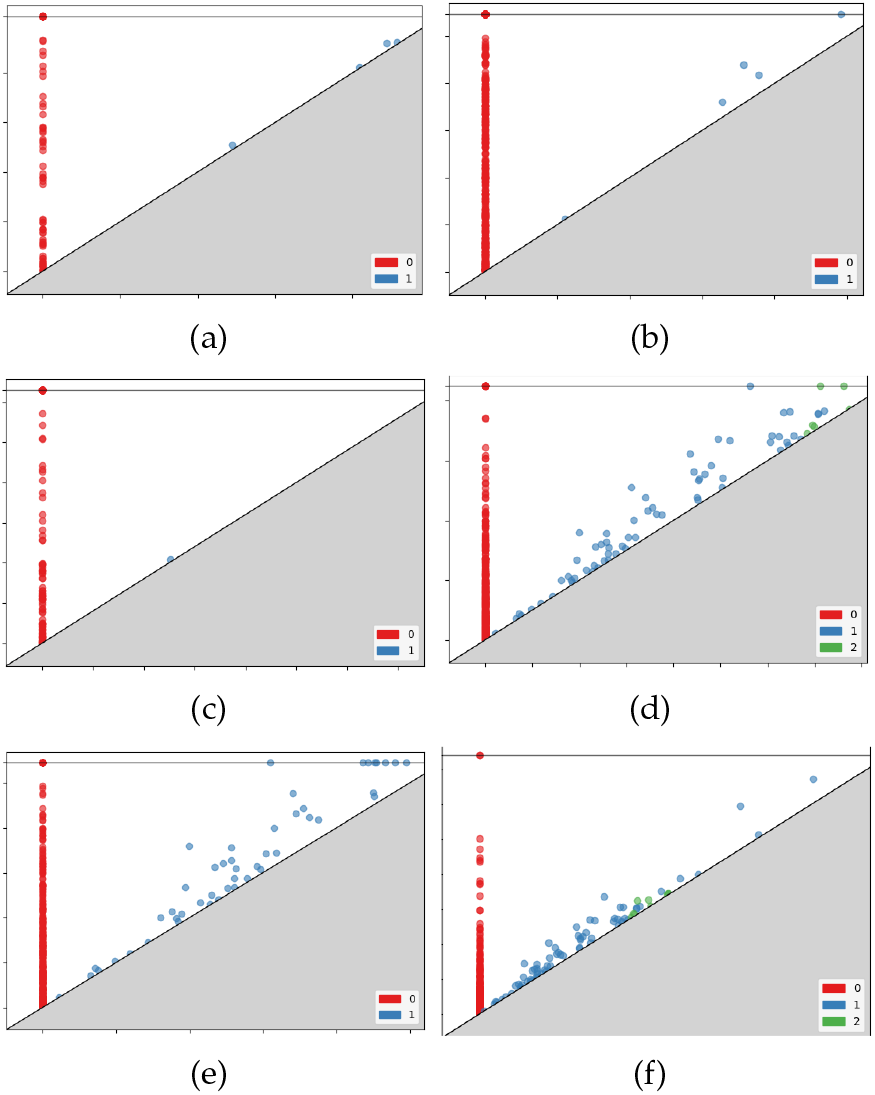
Persistence diagrams for the six clusters of GroEL.

### 3.5 Analysis of Results

Results above establish that the entire methodology of using dimesionality reduction, hierarchical clustering and topological analysis helps in sampling the conformational landscape of a molecule in a way that truly distinct conformations are identified. As mentioned earlier in the section on generation of data, the data for most molecules here was generated using molecular dynamics simulations; they also yield the potential energy of each simulated conformation, taking into account the relative three dimensional placement of each atom that makes the conformation. Topological analysis of the entire energy space is the same as computing persistent homology of the entire embedding generated after feature reduction (which means without filtering the conformations using instance reduction). To explore the energy landscape of the conformational space we filtered the conformations for three molecules (GDP bound CDC-42, Oxytocin and hGalanin, the ones that were generated using MD simulations and were hypothesized to have more than one persistent conformation) based on their energy. We retained only the conformations that have energy less than the 80% of conformations, in other words, we performed filtering at 20% filtration of energy. To do this, we simply sorted the conformations based on their energy and picked the first 20%. The embedding of these molecules at this filtration is shown in Figure-17. As is clear from the embedding, it can be seen that even at such low levels of energy filtration, the molecules separate into two clusters, as suggested by topological analysis. This indicates that the method described in this work to characterize conformational space of proteins is capable of sampling low energy conformations of protein molecules that are likely to persist.

**Fig. 17.**
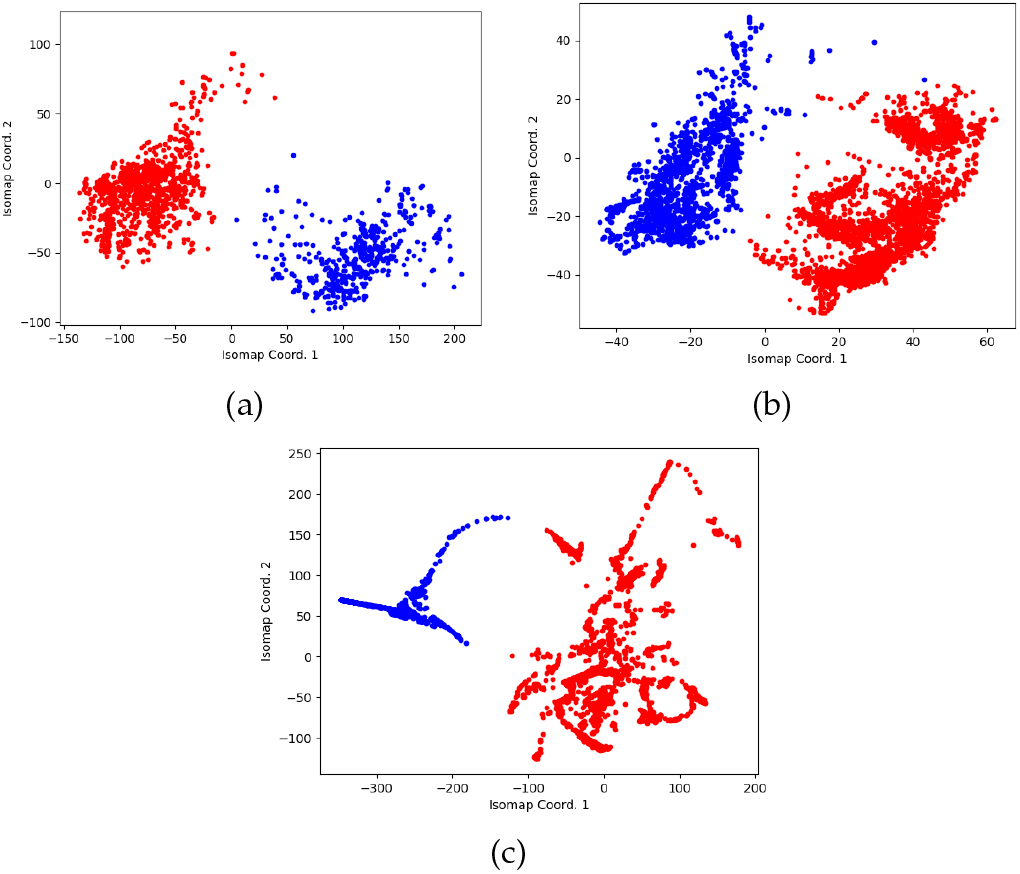
Embeddings of 20% lower energy conformations of: (a)- GDP bound CDC-42 (b)- Oxytocin, (c)- Human Galanin.

## 4 Conclusion

Many proteins undergo large-scale conformational changes as part of their function. Characterizing the conformational space of proteins is crucial for understanding their function and dynamics. We present an efficient filtration procedure that works well for sampling the intermediate conformations for protein molecules that undergo large scale conformational changes as well as for the ones that have a pervasive native state. The method presented is well suited to establish distinctiveness among the clusters generated. Analysis of low energy conformation clusters using dimensionality reduction and algebraic topology and observing its effect with varying energy levels can produce interesting results and help in designing of confomational landscapes for targeted drugs. Refining these methods to produce finer sampling and study the significance of fleeting high energy conformations is the goal ahead. Hierarchical clustering merges (or splits) two sets of conformations based on the heterogeneity of the entire set. In future, we aim at developing a method to bias this divide in a way that is more suited for protein datasets. Molecules that undergo rigorous changes in structure are the ones more suited for such work. In particular, having known intermediates, would help in guiding the conformational pathway search problem as well. It can divide the search space into smaller instances of the same problem. This is also a portion of the ongoing research.

## Footnotes

1. it is predefined, takes care of whether a conformation would be feasible, keeping in mind atom collisions

2. the conformational pathway

## Notes

### Competing Interest Statement

The authors have declared no competing interest.

